# Strategy guides feature recall of a single object in working memory task

**DOI:** 10.1101/2020.05.08.085472

**Authors:** Rakesh Sengupta, Christelle M. lewis

## Abstract

Over past couple of decades our understanding of visual working memory (VWM), and working memory in general, has been predominantly in line with the capacity debate. We recently opened a new line of inquiry regarding the recall of a single object to go beyond the capacity debate, and showed that a series of feature probe questions about a single object yields poorer recall later in the sequence (Sengupta et al, 2020). In the current work we focused on another aspect of sequential feature recall - mainly regarding whether recall can be improved by asking the same question twice. To that end, we chose to focus on two features - color and location, and we contrasted repeat and non-repeat (from the standpoint of feature questions) trials in a series of two experiments. In repeat trials either color or location would be probed twice consecutively. In non-repeat trials color and location probes were presented one after the other in random order. In all trials the stimulus was a small colored oriented line presented for 1 sec in a location within 4o of visual angle. The recall of color and location were mapped onto continuous variable like Sengupta et al, 2020 - for instance, color recall was mapped onto a color wheel. In the first experiment, we used an unaltered color wheel when the color question was repeated. For the second experiment, we used a rotated color wheels for two consecutive color recall trials. We observed an increase in recall error for both repeat and non-repeat condition for location when the probe was at the second question in both experiments. However, color recall error did not increase for second repeat question condition in Experiment 1 as opposed to the non-repeat condition. On the other hand, in Experiment 2 we observed the expected increase in recall error for both repeat and non repeat condition for color probe at the second question. This maybe due to the fact that participants used an ‘anchoring’ strategy in Experiment 1 by remembering where they clicked on the color wheel in the first question. The rotation of color wheel in second experiment destroys the anchor leading to the aforementioned result. The results show that trying to recall the same feature again leads to degradation of recall accuracy for both color and location, and human beings may use different strategies for recall in working memory tasks.

## Introduction

Empirically visual working memory (VWM) has been investigated with two main paradigms - Change Detection, and Serial Recall. Change detection tasks test a person’s ability to encode and recall simultaneously presented items. The standard experiments in this paradigm (Luck and Vogel, 1997; Heyselaar et al., 2011; Vogel and Machizawa, 2004; Woodman et al., 2012; Vogel et al., 2001) have revealed that human performance suffers severely for set sizes more than 4-5. Such results along with similar tests in serial recall domain (Waugh and Norman, 1965; Knops et al., 2014) has led to the idea of visual working memory (VWM) as a limited capacity system (Cowan, 2001).

The main debate in this field for last decade or so has been regarding the exact nature and source of such capacity limitation. In this regard two ideas have often been at loggerheads to explain the behavioral and neuroimaging data in VWM experiments. First one is the ‘slot’ model made popular by (Luck and Vogel, 1997). They endorse the idea of VWM as a limited capacity container for discreet fixed resolution items - or ‘slots’. The other idea held by (Ma et al., 2014) promotes that VWM capacity comes from limited neural resource that has to be distributed dynamically over all items that need memory maintenance.

Both the accounts mentioned above do not take the actual sensory processing into account, and thus quite a lot about what those slots are or what the resources are stretched to cover, is left up to further interpretation and assumption. Moreover treating visual processing like a blackbox leads to an anemic account of VWM, where it becomes difficult to understand the relation between vision, attention, executive control function, and working memory^1^.

In our previous work (Sengupta et al., 2020) we proposed a new paradigm where we tested the recall for a single object probed for multiple features in a serial manner. We showed that every subsequent recall probe reduced accuracy of recall for color, location, and size. For instance, participants were worse at recalling color of the object in memory array if the corresponding question was posed second in the sequence than at first, and so on. We also showed that this recall error was not due to increased temporal delay between subsequent questions. We postulated that this increase in recall error was due to a spontaneous process in response to fulfilling some internal criterion like answering a question. The results posed a challenge to both slots and resource based paradigms, as the memory array populated with only a single object should have ideally all the available slots, or neural resources dedicated to maintaining it in memory. In the current work we investigate whether we can get a better recall by probing the same feature twice, as opposed to asking for different features of the same object in a given trial. In the following we will describe the methods and results.

## Experiment 1

### Apparatus

Participants were invited to do a simple memory test. After being given the instructions on how to do the experiment, they performed the experiment in a dark room (first under guidance for practice and then on their own). They were seated in front of a computer screen with their head stabilized 60 cm away from the LCD screen (34.5cm*19.4cm, resolution:1366*768, refresh rate:60.00Hz). The experiment was designed in Psychtoolbox and was maintained by Octave 4.4.1.

### Stimulus

A Short colored rectangle lines (Width: 5 pixels) of varying length and orientation appeared in various locations from trial to trial. Only one rectangular bar of a random combination of the four features (color, length,orientation and location) was shown per trial. While the stimuli size varied from 1 to 3 degrees of visual angle, the location was set to vary from 0 to 4 degrees of visual angle, and the orientation varied across 360 degrees. The color for each stimuli was randomly extracted from an equi-luminant (*L* = 50) CIELAB color space, the conversion of which is based on Bruce Lindbloom’s website about color spaces.

### Design

After reading the instructions, the experiment began when the subjects were ready. The experiment(s) had 200 trials (each). Each trial began with a small red fixation cross on the center of a white screen for 500 ms. This fixation cross indicated the beginning of every new trial. Immediately after, a short colored rectangular line inclined at a random angle appeared across the screen and remained so for 1000 ms. The stimuli for each trial had random combinations of different colors, lengths, orientations, and locations. The subject was asked to remember (register) the color and the location of the stimuli. After a 1000 ms retention period, the subjects were asked two questions (sequentially) about the stimuli they saw: The color and the location of the stimuli. In some trials, the subject’s were first asked what color the line was and then what the location of the stimuli. In some other trials, they were first asked where the stimuli appeared and then what was the color of the stimuli. In the rest of the trials they were either asked what color the stimulus was twice or where the location of the stimuli was (where the stimulus appeared) twice. The question remained on the screen until the subjects responded. Answers could be reported with a single right mouse click. They judged what the color of the stimulus was by selecting a point on a color wheel consisting of 360 graduations of the primary colors on the circumference of a circle. The colors were chosen from the CIELAB Colorspace (*L* = 50). On the color wheel, the distance between the color of the stimulus shown in that particular trial, and the selected color indicated by the participant, was taken as the error for the color response. To judge the location of the stimulus, a blank screen was shown with the location question presented on the top corner of the screen. The subject had to click at a point they thought was the midpoint of the line (the stimulus). The difference between the selected point and the actual midpoint was taken as the error for the location. A blank white screen was presented for 2000 ms after the two questions (4 combinations) were answered. Each experiment took 20-30 minutes to complete. After the subject completed 25%, 50% and 75% of the trials, they were given a 10-15 minute break that they could take. (They were also allowed to take breaks voluntarily in between the experiment after selecting the second question but not clicking on it.) The experimental design is represented in Fig. 1.

**Figure 1.**
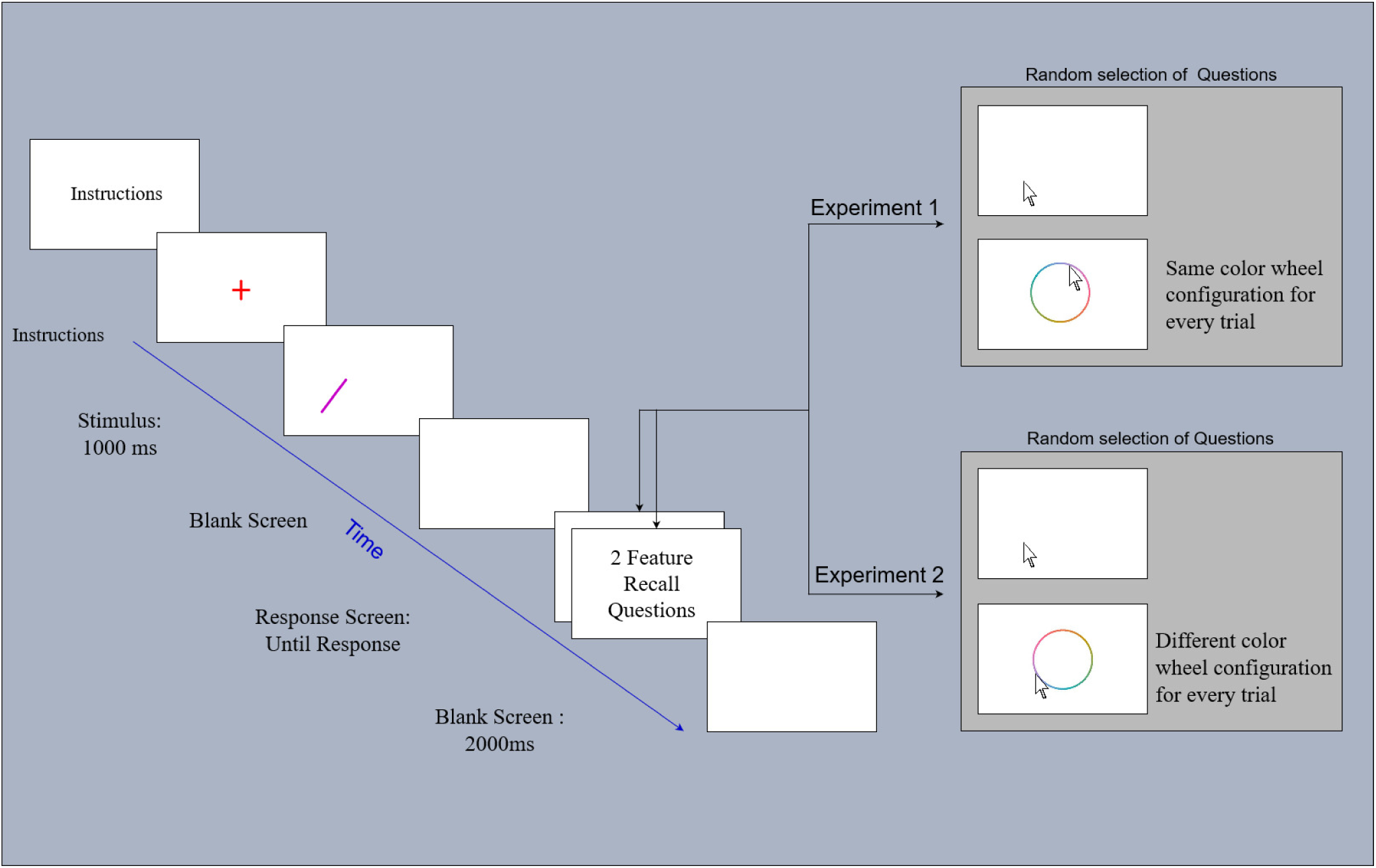
Each trial began with a fixation cross displayed for 500ms. The stimulus following this was a short colored, oriented line at a variable distance away from the center at a duration of ∼1s. After a retention period of ∼1s where a blank screen was displayed, two recall questions were asked: To judge what the color of the stimuli and/or where the stimuli appeared in that trial. Either the color question and the location question was asked in random order or the color question and the location question was repeated. Experiment 1 and 2 had 200 trials each and was both intermittent with breaks. In Experiment 1, the color wheel remained the same across all the trials of the experiment whereas, the orientation of the color wheel varied randomly in Experiment 2.

All subjects were asked two questions in each trial. There were four kinds of conditions. There was *color repeat* and *location repeat* conditions where the color query and location query was repeated, respectively. In the non-repeat condition, we had color and location questions asked in random order - thus we had *color first* and *location first* conditions. When the color question was repeated, the combinations and orientations of colors on the color wheel remained the same throughout the experiment. Thus, in trials where the color question was repeated, the color wheel remained unchanged.

### Participants

19 healthy undergrad students (5 male, 14 female 18 right handed and 1 left handed, with average age of 20 years) with normal to corrected-to-normal vision, and normal color perception, were recruited from SR Engineering College. Informed consent was obtained from all participants before commencing the experiment.

## Results

In line with our previous work (Sengupta et al., 2020), we estimated color recall errors as with angular distance on the color wheel and location recall error as measured in degrees of visual angle. Recall errors for each feature as converted to corresponding *z* scores before analysis. See Table A1 for raw error values.

The results in 2 show that that location recall error increases for the second position in both repeat and non repeat condition. However, color recall seems to not suffer a penalty in the second question in the repeat condition as opposed to non-repeat condition. We conducted a two-way within-subject repeated measures ANOVA, where one factor was the ordinal value of the question in sequence (first or second) and the other factor was the four conditions (color-repeat, color-non-repeat, location-repeat, and location-non-repeat). The results showed a main effect of serial position of feature question (*F*(1, 72) = 148.11, *p* < 0.001, *η*^2^ = 0.45, 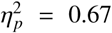) and a significant interaction term (*F*(3, 72) = 36.67, *p* < 0.001, *η*^2^ = 0.33, 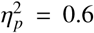). For the post-hoc analysis, we calculated feature-wise Cohen’s d values as effect size between recall errors at different ordinal position for the recall question. For color, the non-repeat condition showed large effect size (Cohen’s d = 1.92, *F*(1, 37) = 42.8, *p* < 0.001) and the repeat condition showed a small effect (Cohen’s d = 0.37, *F*(1, 37) = 1.60, *p* = 0.21). Location recall error showed large effect for both non-repeat (Cohen’d = 3.3, *F*(1, 37) = 126.46, *p* < 0.001) and repeat condition (Cohen’s d = 1.54, *F*(1, 37) = 27.73, *p* < 0.001).

For the repeat condition we saw a very small effect of the position of recall question for color. We hypothesized that this may be due to the fact that color wheel presented during query was the same in both trials. The participants may just be using a strategy where they remember the location on the color wheel they clicked on in the first question and then tried to repeat that in the second question. This can be observed in the increase in standard error for the second question in color repeat condition. Thus we decided to conduct a second experiment where the color wheel will be rotated randomly in every trial, so that no two trials will see the same configuration of the color wheel. All other aspects of design for both experiments remained identical.

## Experiment 2

### Participants

18 healthy undergrad students (11 male and 7female, 17 right handed and 1 left handed, with average age of 20 years) with normal to corrected-to-normal vision, and normal color perception, were recruited from SR Engineering College. Informed consent was obtained from all participants before commencing the experiment.

### Design

A high variance in memory of the color of the stimuli could be a cause of the anchoring effect. To remove what could have been due to an anchoring effect caused by the first color question onto the second color question in the same trial, the color wheel’s orientation was randomized throughout the experiment. Hence, if the color question was repeated in the same trial, the second question would not have an anchor due to the response of the first color question. All the experimental parameters remained the same as Experiment 1.

## Results

The results depicted in Fig. 3 are almost identical to that in Experiment 1 except for one crucial factor. Now the results for both repeat and non-repeat condition are nearly identical with both showing increase in recall error for the second question. The results show a main effect of serial position of feature question (*F*(1, 68) = 95.93, *p* < 0.001, *η*^2^ = 0.42, 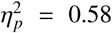) and a significant interaction term (*F*(3, 68) = 20.84, *p* < 0.001, *η*^2^ = 0.28, 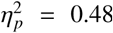). Post-hoc results for color recall show a large effect for both non-repeat (Cohen’s d = 0.89, *F*(1, 35) = 8.77, *p* < 0.01) and repeat conditions (Cohen’s d = 1.0, *F*(1, 35) = 11.19, *p* < 0.01). Location error shows large serial position effect of feature question for both non-repeat (Cohen’s d = 2.49, *F*(1, 35) = 69.05, *p* < 0.001) and repeat conditions (Cohen’s d = 1.8, *F*(1, 35) = 36.05, *p* < 0.001). See Table A2 for raw error values.

**Figure 2.**
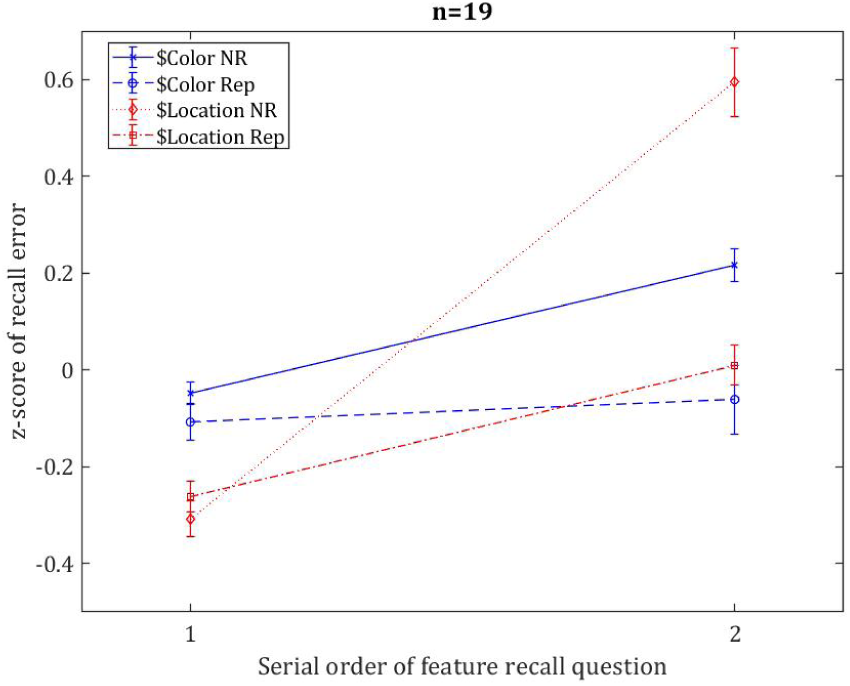
Average *z*-scores across the participants for color and location recall error question in the order of the serial position of the question asked in Experiment 1. In repeat condittion the same feature (color or location) is probed twice. The non-repeat condition is similar to trials in Experiment 2 of Sengupta et al, 2020.

**Figure 3.**
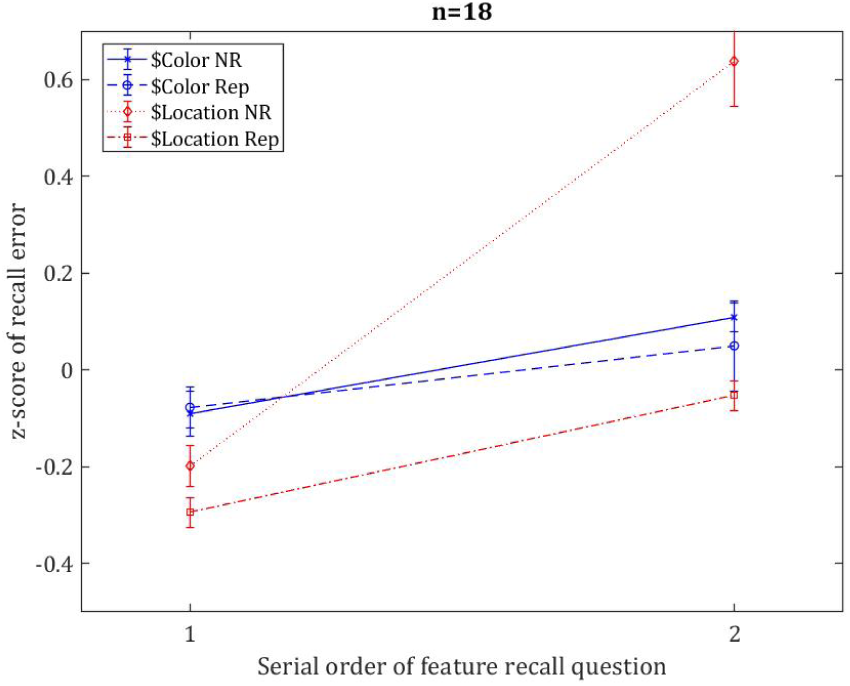
Average *z*-scores across the participants for color and location recall error question in the order of the serial position of the question asked in Experiment 1. In repeat condition the same feature (color or location) is probed twice. The non-repeat condition is similar to trials in Experiment 2 of Sengupta et al, 2020.

## Discussion

The mechanisms of working memory postulated by Baddeley (2003) involve a rehearsal that allows information to be maintained within working memory until completion of the task at hand or intrusion caused by new information provided by attentional mechanisms (Cowan, 2001). In our previous work (Sengupta et al., 2020) we showed that it retention within working memory is more complex and prone to forgetting even when dealing with a single object over the course of 3-5 seconds. We found that a spontaneous erasure of at least a portion of retained memory takes place as soon as participants answer even just one feature recall question, as evidenced by increased recall error in questions answered later in the sequence. This increase in recall error was not prompted by increased delay in subsequent questions as seen in Experiment 3 in Sengupta et al. (2020).

In the current work we intended to check whether participants can reduce recall error by answering the same question twice. We chose to focus on two features - color and location. We had repeat and non-repeat conditions. In repeat condition the same feature recall question was repeated twice, in non-repeat condition color and location questions were asked in random order (see Fig. 1). In Experiment 1, the color wheel presented at the corresponding question stage was of the same during the two questions in color repeat condition. In Experiment 2 the configuration of the color wheel was rotated by a random angle in each trial. We see from Fig. 2 and 3 that location recall errors increase at the second question regardless whether it was in repeat of non-repeat trial in both Experiment 1 and 2. The magnitude of the recall error was slightly smaller in the repeat condition for location in both experiments. However, in Experiment 1 color recall error did not suffer a penalty in the repeat condition as opposed to non-repeat condition. The situation changed in Experiment 2 where color recall error increased nearly identically for the second serial recall question position for both repeat and nonrepeat trials.

The results point to a very interesting strategy that participants employed for color recall in Experiment 1 in repeat condition trials. The participants while answering the color question tried to remember where they clicked on the color wheel and tried to repeat the same when asked about color second time, as seen in the increased variance of recall error observed in the second trial in repeat condition (see Fig. 2). The strategy was rendered ineffective in Experiment 2 where we randomized the configuration of the color wheel in every trial. This points to two very important conclusions we can draw from above. Firstly, the increase in recall error we observed in Sengupta et al. (2020) can not be removed by simply repeating the question again. Rather it seems that it is surely permanent loss of memory precision that can not be reconstructed without other strategies. Secondly, participants can employ strategy unrelated to the feature in question in order to increase performance in working memory task as seen in Experiment 1.

Combined together, the results lead us to suggest that we need to revisit the mechanisms of working memory and possibly include behavioral strategies (visual or non-visual) that cognitive agents can use in order to improve VWM performance. Further work is needed to explore possible neural correlates of this phenomenon.

## Appendix

### Raw results

**Table A1.**
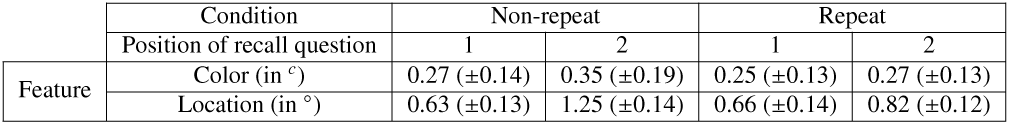
Raw recall errors for all the conditions in Experiment 1. Color recall errors are expressed in radians for the angular difference on color heels. Location errors are expressed in degrees of visual angles. Standard errors of mean are reported in parenthesis.

**Table A2.**
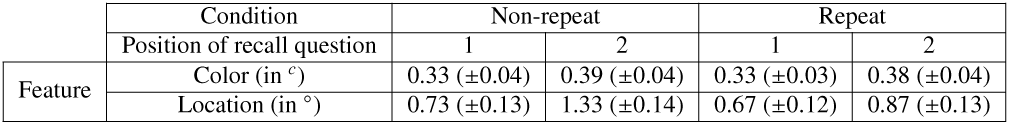
Raw recall errors for all the conditions in Experiment 2. Color recall errors are expressed in radians for the angular difference on color heels. Location errors are expressed in degrees of visual angles. Standard errors of mean are reported in parenthesis.

## Footnotes

1 The blackbox approach has led to a limited scope of behavioral research as well. With fewer parameters to manipulate, psychophysical tests of VWM has led to experimenters researching a very narrow part of the visual field. For instance, change detection protocols have mostly adhered to narrow display size (within 10° of visual field around fovea (Luck and Vogel, 1997; Heyselaar et al., 2011; Vogel and Machizawa, 2004; Woodman et al., 2012; Vogel et al., 2001). In doing so we are limiting the scope and understanding of human vision and memory in general.

